# A deep convolutional neural network approach for astrocyte detection

**DOI:** 10.1101/241505

**Authors:** Ilida Suleymanova, Tamas Balassa, Sushil Tripathi, Csaba Molnar, Mart Saarma, Yulia Sidorova, Peter Horvath

**Affiliations:** Laboratory of Molecular Neuroscience, Research Program in Developmental Biology, Institute of Biotechnology (HiLIFE), University of Helsinki, Viikinkaari 5D, FI-00014 Helsinki, Finland.; Synthetic and Systems Biology Unit, Hungarian Academy of Sciences, Biological Research Centre (BRC), Temesvári körút 62, 6726 Szeged, Hungary.; Research Program Unit, Helsinki Institute of Life Science (HiLIFE), Faculty of Medicine, University of Helsinki, Haartmaninkatu 8, 00014 Helsinki, Finland.; Institute for Molecular Medicine Finland (HiLIFE), University of Helsinki, Tukholmankatu 8, 00014 Helsinki, Finland.

## Abstract

Astrocytes are involved in brain pathologies such as trauma or stroke, neurodegenerative disorders like Alzheimer’s and Parkinson’s disease, chronic pain, and many others. Determining cell density and timing of morphological and biochemical changes is important for a proper understanding of the role of astrocytes in physiological and pathological conditions. One of the most important of such analyses is astrocytes count within a complex tissue environment in microscopy images. The most widely used approaches for the quantification of microscopy images data are either manual stereological cell counting or semi-automatic segmentation techniques. Detecting astrocytes automatically is a highly challenging computational task, for which we currently lack efficient image analysis tools. In this study, we developed a fast and fully automated software that assesses the number of astrocytes using Deep Convolutional Neural Networks (DCNN). The method highly outperforms state-of-the-art image analysis and machine learning methods and provides detection accuracy and precision comparable to that of human experts. Additionally, the runtime of cell detection is significantly less than other three analyzed computational methods, and it is faster than human observers by orders of magnitude. We applied DCNN-based method to examine the number of astrocytes in different brain regions of rats with opioid-induced hyperalgesia/tolerance (OIH/OIT) as morphine tolerance is believed to activate glial cells in the brain. We observed strong positive correlation between manual cell detection and DCNN-based analysis method for counting astrocytes in the brains of experimental animals.

## 1. Introduction

Astrocytes are a type of glial cells within the central nervous system (CNS). On average, there are 21–26 billion neurons and 27–40 billion glial cells in a human brain^1^. The proportion of astrocytes in the CNS varies by brain and spinal cord regions and it is estimated to range from 20% to 40% of all glia^2^. Astrocytes respond to injury and CNS diseases, and play an important role in development and maintenance of chronic pain and certain psychiatric disorders, such as autism spectrum disorders and schizophrenia^3,4^.

Morphologically, astrocytes are star-shaped structures with small rounded bodies, and numerous long, ramified branches. These cells are disseminated distinctly over the nervous tissue. Investigations of morphological changes of astrocytes in pathological conditions or after chemical/genetic perturbations is usually carried out through the rigorous process of cell staining, imaging, image analysis and quantification. The versatility of the branching structure of astrocytes makes their automatic detection rather challenging. Therefore, developing fully automated counting methods of immunohistochemically stained images that require no user intervention is a major issue. Traditionally, manual or semi-automated techniques have been used to evaluate the number of CNS cells in samples of interest. However, manual counting processes are time-intensive, cumbersome and prone to human errors. Commonly used open-source tools for general cell quantification such as ImageJ^5^, custom scripts or Ilastik^6^ have technical limitations concerning the accurate detection of astrocytes because of their complex morphology^2,7,8^. Usually they are based on either machine-learning methods (Ilastik) or thresholding (ImageJ, custom scripts)

Currently, no efficient image analysis tools are available to quantify astrocytes from large-scale histology datasets. A novel potential way to circumvent manual or semi-automated techniques is to use Deep Convolutional Neural Networks (DCNN) approaches to identify cells. Deep learning is a type of machine learning approach based on learning from multiple layers of feature extraction, and can be used to analyze complex data such as images, sound and text^9–14^. Recently, DCNN have gained increasing attention in the field of computational cell biology^15–18^ and have shown great success especially in complex image classification tasks^19^.

Here, we propose a DCNN-based method and an open-source software platform enabling biologists and pathologists to accurately detect astrocytes in immunohistological images (**Supplementary Software 1**). Our software, *FindMyCells* (www.findmycells.org), is a fully automated and user-friendly software for more precise detection of cells on microscopy images. We remark, however, that the software can also be used for a range of other object detection tasks if underlying DCNN module is properly trained to recognize these objects. DCNN show tremendous improvement in accuracy in case of detecting complex morphologies such as astrocytes. According to our measurements, FindMyCells strongly outperformed other computational methods and was comparable to the performance of human experts. Very importantly, showed better performance than what we measured between field experts.

To validate the proposed software, we compared the number of cells counted by human experts, and by FindMyCells, as well as by three other software (Ilastik, custom threshold-based script^20^, ImageJ) analyzing brain tissues of rats treated with repeated injections of morphine to induce opioid-induced hyperalgesia/tolerance (OIH/OIT). OIH/OIT is a common complication of prolonged opioid therapy which is characterized by enhanced pain/lack of pain relief^21^. Recent evidences support the role of glial activation in the spinal cord in the development and maintenance of OIH/OIT^22,23^, but the implication of brain glia in this process required further studies. Thus, we analyzed glial cells in different brain regions believed to be involved in generation and maintenance of pain^24,25^ using immunohistochemistry (IHC)^26^. The model, biological experiments and outcomes are described in details by Jokinen and co-authors^26^. In the current study, we first counted the number of astrocytes by FindMyCell, Ilastik, custom threshold-based script^20^ and ImageJ in randomly selected sections from the brains of control and morphine-treated animals described by Jokinen^26^. Thus, we compared the performance of FindMyCells to semi-automated tools based on both machine-learning principles and manual thresholding. To assess applicability of FindMyCells to real laboratory tasks, we selected images from striatum where we previously observed changes in the number of astrocytes^26^ and analyzed them by manual counting and FindMyCells. A very strong correlation was shown between FindMyCell output and manual counting of astrocytes.

## 2. Materials and methods

### 2.1 Preparation of samples and datasets for analysis

An experimental dataset containing microscopy images of several different regions of rat brain was used to train and examine the performance of the proposed DCNN approach. Samples were treated as described by Jokinen and co-authors^26^. Briefly, brains of rats with and without OIH/OIT were sectioned and astrocytes were labelled with glial fibrilar acidic protein (GFAP) antibodies (Cat#G-3893, Sigma-Aldrich, St. Louis, MO, USA), followed by VECTASTAIN ABC HRP Kit (Cat#PK-4002 Vector Laboratories, Burlingame, CA, USA). Slides were scanned using the 3DHISTECH Scanner (3DHISTECH Ltd, Budapest, Hungary). The images for analysis were saved in tiff format from the Pannoramic Viewer software using a 20x magnification. Two datasets were prepared: training data set contained 1200 images from random regions of rat brains and validation data set containing 12 images from striatum. The images from the training dataset were handled as shown in Figure 1. Briefly, after sample preparation, astrocytes were annotated and a deep neural network was trained. Next, DCNN prediction was executed, and cell count and coordinates were collected. Details of these procedures are described below.

**Figure 1.**
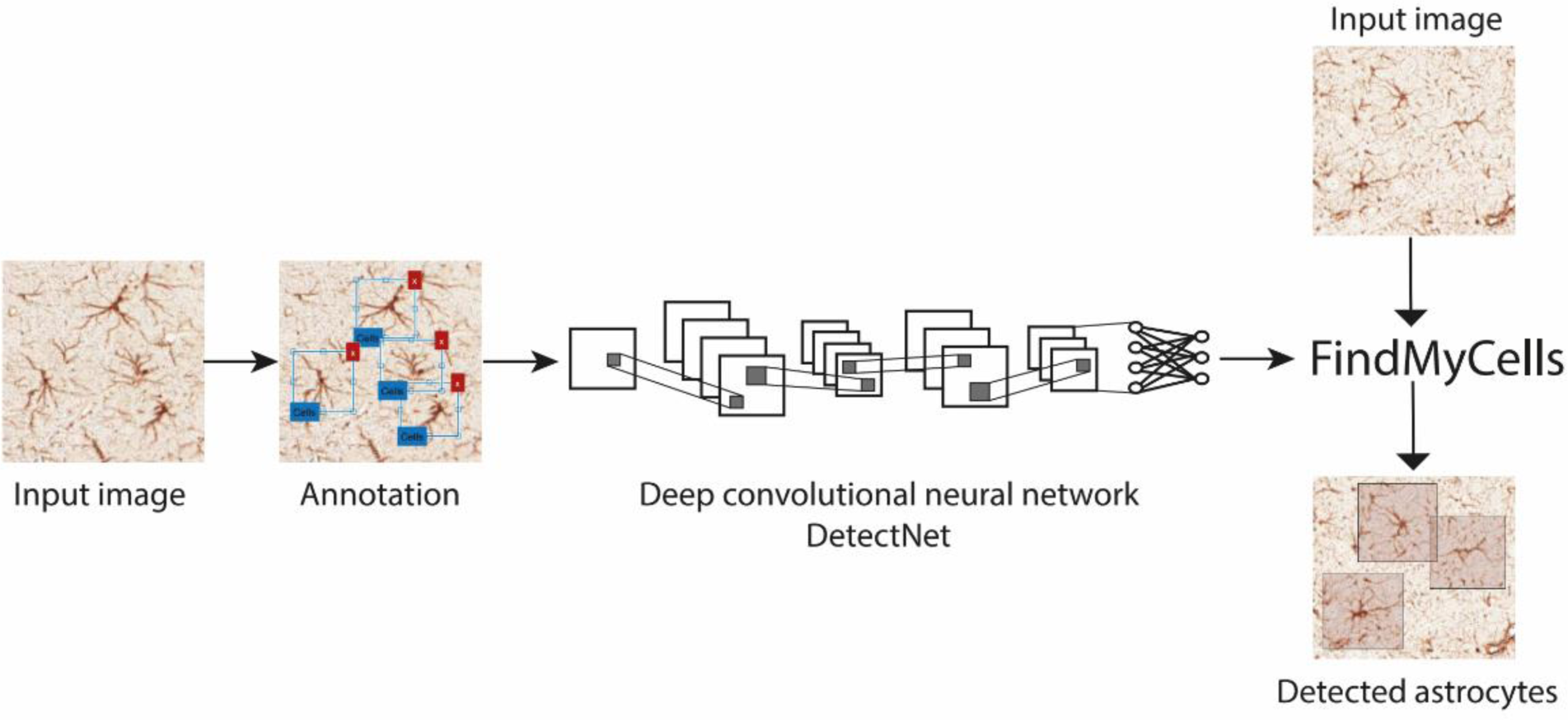
Schematic overview of the proposed framework and the FindMyCells software based on DCNN. Left: 15,000 cells were annotated with their bounding boxes and a DCNN method was trained based on the detectNet architecture for object detection^28^. Right: The FindMyCells software starts from the raw image data, a DCNN model receives the raw images as input and after prediction marks identified astrocytes with bounding boxes around cell body. Software also exports the coordinates of detected cells as a table (not shown on the scheme).

### 2.2 Annotation procedure

Annotations were made manually using the Matlab region of interest (ROI) Training Image Labeler application. The main training set, a total of 1200 images and 15,000 cells was labelled by a single expert, providing a rich platform for testing the accuracy of astrocyte detection (**Supplementary data 1** and BBBC0xx^a^). For performance evaluation, an additional expert was asked to label a smaller test set of ~30 different images containing ~300 cells. This validation dataset was annotated twice at two different time points by both of the experts to measure intra person accuracies.

### 2.3 Software platform and convolutional neural networks

The proposed new tool for astrocyte detection was implemented in Python 2.7+. We developed the software’s graphical interface using the PyQt5 package. For the deep learning implementation, we used detectNet^b^ architecture which is an extension of the *caffe* package^27^. NVIDIA DIGITS^c^ framework was used to train the network. Training computations were conducted on a notebook PC with a 2.3 GHz Intel Core i5, 8GB RAM memory and an NVIDIA GeForce GTX. The software was then run on a 3.5 GHz Intel Core i7–4770K computer with 16 GB memory and an NVIDIA TitanXp graphic card.

### 2.4 Evaluation methods and metrics

To evaluate the performance of the proposed framework we measured precision, recall and detection accuracies achieved by two different annotators and four different computational methods. For the quantitative characterization of these parameters we made an object matching between the ground truth objects and those of the detection of interest. For matching objects, we used the Hungarian method that provides optimal pairing between point sets. To calculate this matching, we defined pairing as an assignment problem. We assigned a weight to each pair of ground truth objects and detected matching objects. The assigned weight was set to infinite if the objects in a pair had no overlap. Otherwise the weight was set to the reciprocal of the area of overlap. The total cost of a pairing was defined as the sum of all the weights of paired objects^29^.

A correct detection (true positive, TP) was established when a ground truth object had a matched pair. False positive (FP) detection was quantified when an extra object was present on the applied method’s output, while a false negative (FN) was considered in case of missing objects. Based on these; precision (P) = TP/(TP+FP), recall (R) = TP/(TP+FN), and detection accuracy (DA) = TP/(TP+FP+FN) were calculated.

We compared a total of six sets of astrocyte detection data, of which two were generated by human experts, one by FindMyCells pipeline, one by Ilastik, one by thresholding and simple morphology operations^29^, and one by using standard ImageJ operations. To measure the self- and cross-accuracies of human experts the same test images were blindly shown to the annotators for re-labelling. To minimize the bias, the duplicate images were mirrored and/or rotated. Altogether ~30 different images containing ~300 cells were manually annotated using the Matlab ROI Training Image Labeler application.

In Ilastik, we used pixel and object classifications with Gaussian Smoothing color/intensity feature, Gaussian Gradient Magnitude and Difference of Gaussians edge features. For the threshold based method we chose an algorithm with manual thresholding offered by the Matlab software. The algorithm detects positive cells when the intensity of their staining is substantially higher than the background. All pixels outside the region of interest are set to zero. Matlab’s built-in functions trace boundaries cells/objects. Objects with the size smaller than the average size of cells were excluded from the quantification using object size threshold set manually^20^. For ImageJ, we used intermodes threshold, has a substantial gain in removing background noise and identifying individual cell in the image, objects with the size smaller than the average size of cells were also excluded as for the threshold based method.

## 3. Results

We compared the performance of the two field experts and of the four computational methods for the quantification of astrocytes using precision, recall, and detection accuracy metrics (Figure 2) calculated as described in methods section. Figure 2a shows examples of astrocytes detection by FindMyCell software on images.

**Figure 2.**
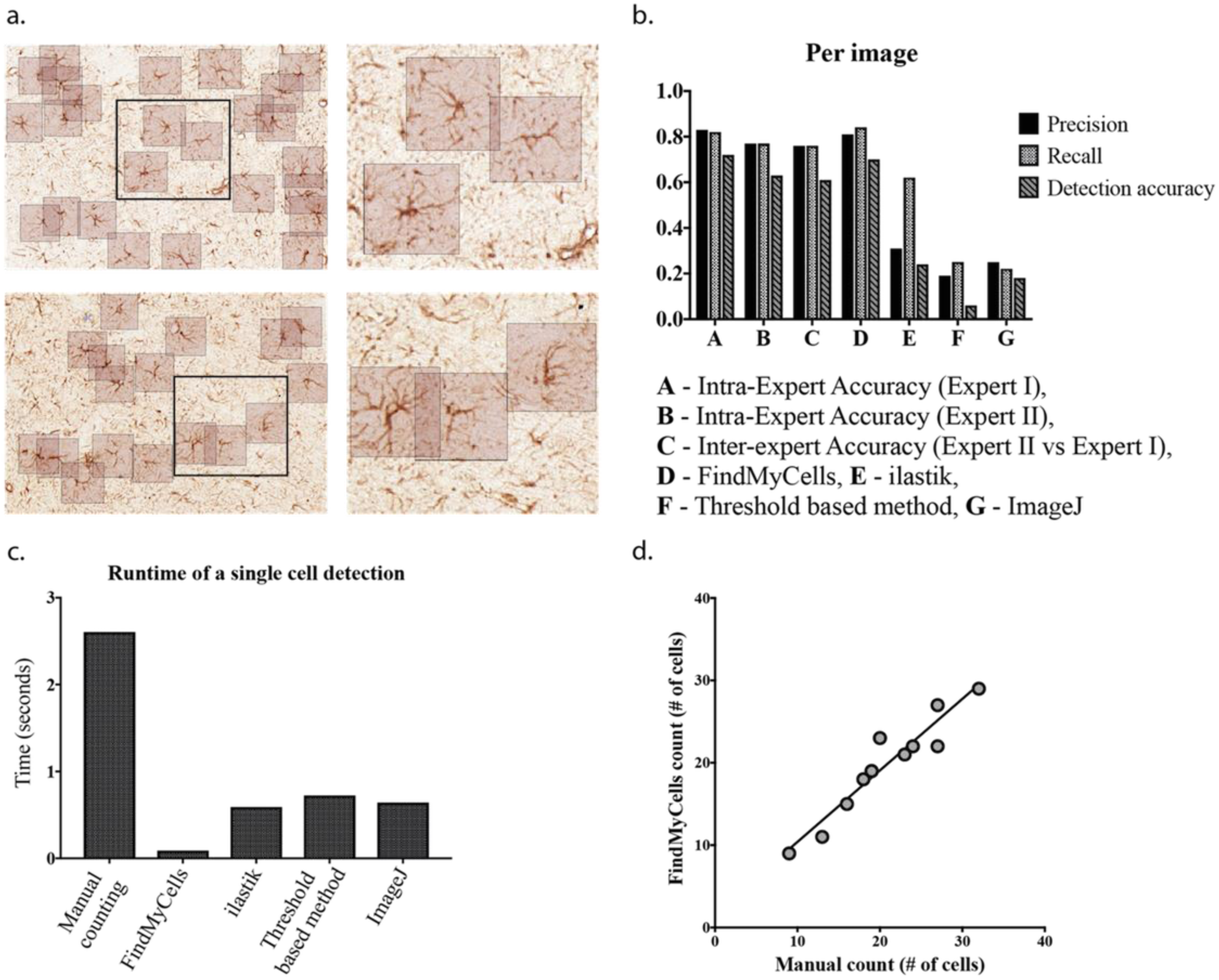
Examples of astrocyte detection by FindMyCells. **a**: detection results. **b**: precision, recall and detection accuracy values for the human experts, FindMyCells, Ilastik and a threshold-based algorithm. **c:** average detection time of human observer and the computational methods. **d**: Per image Pearson correlation between FindMyCells output data and manual counting of astrocytes.

Figure 2b illustrates the comparison of the results of application of different quantification methods to image analysis. First, we measured self-accuracy of the human experts. As expected, astrocyte detection proved to be a complex issue, thus even intra-observer accuracy was far from being perfect. Detection accuracy was 0.72 and 0.63 for the two human experts, respectively. We also evaluated inter-observer accuracy between the two human experts for which the detection accuracy was 0.61. Each case the test set made by the first human annotator was used as ground truth, we remark that this same annotator labeled the 15,000 training examples.

We expected that an appropriate computational method would perform at least as good as suggested by the inter-observer accuracy, and might approach the self-accuracy of our experts. Indeed, an outstanding performance was observed for the proposed DCNN. Its detection accuracy, precision and recall values were close to or even better than the human experts’ self-accuracy (P = 0.81 R=0.84, DA = 0.70). We also tested Ilastik (P = 0.31, R = 0.62, DA = 0.24), the threshold-based analysis method (P = 0.19, R= 0.25, DA = 0.06), and ImageJ (P = 0.25, R = 0.22, DA = 0.18) for these parameters. These tools were highly outperformed by the human experts as well as by FindMyCell software.

In practice, the runtime of any computational method is an important parameter concerning its performance. Therefore, we evaluated average runtime of different methods to annotate a single cell. For the human experts, it took approximately 2.6 seconds to annotate a single cell. This value is approximately 0.088 seconds for FindMyCells, 0.59 seconds for Ilastik, about 0.72 seconds for the threshold-based method and 0.64 seconds for ImageJ on the same computer (Fig 2c), indicating that FindMyCells is characterized by a highly beneficial runtime besides its good-quality performance. Important to remark that FindMyCells uses highly parallelized GPU calculations. A drawback of FindMyCells is that it required 15,000 annotated cells (~11 hours of continuous labeling) and a strong GPU for training the model (~18 hours of training time, 300 epochs). Statistics for all evaluated metrics are shown in **Supplementary Table 1**.

Finally, we evaluated the proposed method on detecting astrocytes in a control and a drug-treated tissue environment, and compared its detection accuracy with that of the human experts. For this, 12 striatum sections of morphine treated and control animals (6 per group) were used. Cells were counted manually and also by using FindMyCells, respectively, and the cell count ratio for treated groups vs. control was computed. The computed ratio was 0.703 for manual counting and 0.715 for FindMyCells, suggesting that our approach is precise as well as robust in counting cells indicating an exceedingly high, ~99% measurement similarity. The proposed method produced coherent results as obtained by the manual counting method at each analyzed image, with a Pearson correlation of R=0.95 (Fig 2d).

## 4. Discussion

In this paper we introduced a novel, fully automated software FindMyCells (www.findmycells.org) for accurate detection of astrocytes, that is need to better understand the role of these cells in pathophysiological processes occuring in CNS as a result of trauma, stroke, neurodegenerative disorders or chronic pain. Astrocytes represent special cell type characterized by a huge variation in appearance, size, and shape that complicates their analysis.

Manual counting is considered to be the gold standard method for astrocyte detection and it is the most widely^30^ used approach in the world^31–32^. However, manual approach is prone to the operator’s subjectivity, and is associated with non-reproducible and imprecise experimental results. Furthermore, subjective bias may occur when the operator is not blinded to the sample.

We propose a DCNN-based method and an open-source software platform enabling biologists and pathologists to accurately detect astrocytes in immunohistological images. FindMyCells is a fully automated and user-friendly software for the precise detection of these cells in bright-field images. To train the DCNN module in our software for this paper we labelled 15,000 cells manually and achieved detection accuracy of 84%. The detection accuracy of FindMyCells software might be further improved when more annotations are available. Nevertheless, even in the current state FindMyCells significantly outperforms tested semi-automated methods based on both machine-learning principles (Ilastik) and classical image analysis, and has detection accuracy comparable to that of human experts. Importantly, we observed pronounced variation in detection accuracy between two human experts, although both of them came from the same lab with similar domain knowledge. We consider that the fairly low level of intra-observer accuracy indicates the challenging nature of reliably detecting astrocytes.

To validate the proposed system, we compared the human observer’s detection-based count with the performance of FindMyCells. Our results on real data show that upon measuring the ratio of astrocytes in the brains of morphine treated and control rats FindMyCells output data differ from counts produced by the human expert by only 1%, making the proposed method highly applicable for real tasks in everyday practice. Importantly, these results were in line with previously published data^26^.

In addition to outstanding detection accuracy, runtime of FindMyCells is really low. The software requires approximately 30-fold less time to detect single cell on the image than human expert. We remark here that our DCNN implementation efficiently utilizes parallel GPU capabilities. As a drawback, the annotation of the cells on images and training of DCNN module is time consuming.

Our future plans include the extension of FindMyCells with a comfortable web-based annotation and a training module, and also with a server system where users with limited GPU access can submit their annotated dataset to our GPU cluster for the training of their custom DCNN. If it is supported by community efforts we will hopefully collect versatile annotated data and develop universal software for detection of various cell types.

## Author contributions

P.H. conceived and led the project. M.S and Y.S. co-supervised the project. T.B. designed and developed the software. I.S. wrote the documentation, collected the data and prepared the figures. P.H., I.S., T.B., and M.S. designed the experiments and analyzed the data. S.T., I.S., T.B., M.S., Y.S., and P.H. wrote the manuscript. All authors read and approved the final manuscript.

## Acknowledgments

P.H. acknowledges support from the Finnish TEKES FiDiPro Fellow Grant 40294/13, IS, YS and MS acknowledge the support from the EU FP7/2007 - 2013 under grant agreement No 602919. B.T., C.M., and P.H. acknowledge funding from the European Union and the European Regional Development Funds (GINOP-2.3.2-15-2016-00006, GINOP-2.3.2-15-2016-00026). The authors would like to thank Prof. Eija Kalso and Prof. Perka Rauhala for support, Dr. Viljami Jokinen and Dr. Hanna Viisanen-Kuopila for sharing samples/images used in our study, Jenni Montonen for assistance with cells annotation and data collection, Dora Bokor, PharmD, for proofreading the manuscript.

## Footnotes

a The annotations and the image data was deposited to the Broad Bioimage Benchmark Collection

b https://devblogs.nvidia.com/parallelforall/detectnet-deep-neural-network-object-detection-digits/

c https://github.com/NVIDIA/DIGITS

